# scCaT: an explainable capsulating architecture for sepsis diagnosis transferring from single-cell RNA sequencing

**DOI:** 10.1101/2024.04.17.590014

**Authors:** Xubin Zheng, Dian Meng, Duo Chen, Wan-Ki Wong, Ka-Ho To, Lei Zhu, JiaFei Wu, Yining Liang, Kwong-Sak Leung, Man-Hon Wong, Lixin Cheng

## Abstract

Sepsis is a life-threatening condition characterized by an exaggerated immune response to pathogens, leading to organ damage and high mortality rates in the intensive care unit. Although deep learning has achieved impressive performance on prediction and classification tasks in medicine, it requires large amounts of data and lacks explainability, which hinder its application to sepsis diagnosis. We introduce a deep learning framework, called scCaT, which blends the capsulating architecture with Transformer to develop a sepsis diagnostic model using single-cell RNA sequencing data and transfers it to bulk RNA data. The capsulating architecture effectively groups genes into capsules based on biological functions, which provides explainability in encoding gene expressions. The Transformer serves as a decoder to classify sepsis patients and controls. Our model achieves high accuracy with an AUROC of 0.93 on the single-cell test set and an average AUROC of 0.98 on seven bulk RNA cohorts. Additionally, the capsules can recognize different cell types and distinguish sepsis from control samples based on their biological pathways. This study presents a novel approach for learning gene modules and transferring the model to other data types, offering potential benefits in diagnosing rare diseases with limited subjects.

**Author summary:** Deep learning models used in disease diagnosis usually suffer from insufficient data for training and the lack of explainability, especially in rare diseases. These shortages hinder their application to sepsis diagnosis. Here we propose a diagnostic framework name scCaT(https://github.com/Kimxbzheng/CaT), which transfers knowledge learned from single-cell RNA-seq, for diseases with insufficient bulk data. The framework uses capsulating architecture to group genes into capsules and provide explainability to the deep learning model for sepsis diagnosis. ScCaT achieves robust and outstanding performance for sepsis diagnosis in both scRNA-seq and bulk RNA datasets. This architecture offers potential approaches in diagnosing rare diseases with limited subjects with explainability.

## Introduction

Sepsis is a severe condition with the highest mortality rate in the intensive care unit (ICU), caused by an overreaction of the immune system to pathogens which can damage multiple organs [1]. However, diagnosing sepsis is challenging as its symptoms can also be caused by other disorders, and there are no approved biomarkers for its precision diagnosis. Current diagnosis relies on a series of tests, including blood test, urine test, X-ray and computed tomography (CT)-scan, which is time-consuming and may miss the optimal window for intervention. Therefore, an effective molecular diagnostic model is critical to assist in sepsis detection.

The rapid development of the high-throughput sequencing technologies and the deep learning techniques has led to numerous computational methods for disease detection and diagnosis [2–6]. Recently, large deep learning models such as ChatGPT [7] and GPT-4 [8] have shown impressive performance. Deep learning-based diagnostic models, such as those proposed by Bilal et al.[9], Ethan et al.[10], and Kam et al. [11] have utilized convolutional neural network and long short-term memory for sepsis detection. However, these models rely on vital signs and signals like electrocardiogram (ECG) and biochemistry laboratory values measured by different clinical tests, which could be time-consuming. On the other hand, biomarkers based on gene expression, such as sNIP [12], SeptiCyte [13], and FAIM3/PLAC8 [14], only require a blood test. Nevertheless, they are merely developed based on microarray gene expression data and not generalizable across platforms.

Although deep learning models provide opportunities for molecular diagnostics of sepsis via analysis of host transcriptome data, we face two major challenges. Firstly, training these models requires a substantial amount of data, typically on the order of tens of thousands. In clinical settings, however, it is difficult to acquire such a large dataset due to factors such as privacy concerns, resource-intensive processes, time constraints, and limited participant availability, particularly for rare diseases or complex medical conditions. Secondly, the deep learning models inherently lack explainability, often functioning as “black boxes” with little insight into the underlying biological mechanisms driving their predictions. This lack of interpretability poses challenges for clinicians and researchers in understanding the decision-making process of the models.

To overcome these challenges, we proposed a deep learning framework called scCaT that fused capsulating architecture and Transformer to develop a detection model for sepsis diagnosis using single-cell RNA sequencing (scRNA-seq) data. ScRNA-seq provides high-resolution gene expression data at individual cell level, but the model trained solely on scRNA-seq data is not readily adapted to bulk RNA measurements, which are commonly used in clinical diagnostics. To address this, transfer learning was applied to transfer the model to bulk RNA cohorts. Specifically, the model trained on the scRNA-seq dataset was utilized as a pre-trained model and was fine-tuned on microarray and bulk RNA-seq data.

Moreover, scCaT utilized the capsulating architecture as an effective gene expression encoder to improve explainability. We functionally characterized the capsulating architecture, also referred as capsule network, which groups genes with similar biological functions into capsules by dynamic routing owing to genes tend to work together as modules during the biological process. Subsequently, the self-attention mechanism known as Transformer was employed as a classifier to identify the cells from sepsis patients or controls.

ScCaT achieved high accuracy with an area under the receiver operating characteristic curve (AUROC) of 0.93 on single-cell test set and an average AUROC of 0.99 on six microarray independent cohorts. Moreover, the results demonstrated that the capsule network learned to recognize different cell types and distinguished sepsis in each cell type based on the biological functions grouped in 20 capsules. The 20 capsules were found to be enriched in the immune-related functions in the inflammation associated with sepsis.

By combining the strengths of the capsulating architecture, Transformer, and transfer learning, scCaT addresses the problems of data availability, adaptability to other data types, and the need for applying explainability in deep learning models for disease diagnosis. Our framework shows promising potential for improving the accuracy and interpretability of sepsis detection, and it paves the way for future advancements in the diagnosis and the prediction of outcomes for diseases and medical conditions with limited data.

## Material And Methods

### Study Design

We collected single-cell RNA sequencing (scRNA-seq) data for septic patients and normal controls from the Broad Institute Single Cell Portal (https://singlecell.broadinstitute.org/single_cell), with portal ID: SCP548 (subject PBMCs). The dataset contains gene expression data for 126,351 cells from 29 septic patients and 36 corresponding controls across three cohorts. The septic patients include subjects with urinary-tract infection and mild or transient organ dysfunction (Int-URO), clear or persistent organ dysfunction (Urosepsis, URO), and sepsis in hospital wards (Bac-SEP) and medical intensive care unit (ICU-SEP). The control samples included subjects with urinary-tract infection but no organ dysfunction (Leuk-UTI), subjects admitted in the medical intensive care unit without sepsis (ICU-NoSEP), and uninfected healthy controls. We filtered out cells that profiled less than 20% of genes and genes that were recorded in less than 20% of cells.

We also collected data from nine microarray cohorts that profiled the blood of septic patients and normal controls (**Table 1**). We then intersected the genes profiled in SCP548 with those in RNA-seq and microarray cohorts, resulting in the expression data for 2,869 genes used for downstream analysis.

**Table 1.** Single-cell RNA-seq and bulk RNA cohorts used in this study.

| Dataset accession | Data type | Sample type | Number<br>of septic<br>subjects | Number<br>of normal<br>subjects | Total<br>subjects | Reference |
| --- | --- | --- | --- | --- | --- | --- |
| Discovery cohort |  |  |  |  |  |  |
| SCP548 | Single-cell<br>RNA-seq | PBMC | 29 | 36 | 65 | Ref. [26] |
| Fine-tuned cohorts |  |  |  |  |  |  |
| GSE185263 (30%) | RNA-seq | Whole blood | 103 | 14 | 117 | Ref. [27] |
| GSE95233 | Microarray | Whole blood | 102 | 22 | 124 | Ref. [28] |
| GSE26440 | Microarray | Whole blood | 98 | 32 | 130 | Ref. [29] |
| GSE57065 | Microarray | Whole blood | 82 | 25 | 107 | Ref. [30] |
| Validation cohorts |  |  |  |  |  |  |
| GSE185263 (70%) | RNA-seq | Whole blood | 245 | 30 | 275 | Ref. [27] |
| GSE28750 | Microarray | Whole blood | 10 | 20 | 30 | Ref. [31] |
| GSE8121 | Microarray | Whole blood | 60 | 15 | 75 | Ref. [32] |
| GSE9692 | Microarray | Whole blood | 30 | 15 | 45 | Ref. [33] |
| GSE13904 | Microarray | Whole blood | 52 | 18 | 70 | Ref. [34] |
| GSE26378 | Microarray | Whole blood | 82 | 21 | 103 | Ref. [29] |
| GSE4607 | Microarray | Whole blood | 69 | 15 | 84 | Ref. [35] |

### ScCaT

The proposed deep learning framework, scCaT, incorporated capsulating architecture, capsule network, and the self-attention model, Transformer. The capsule network served as an encoder that encoded effective genes’ expression into capsules, which can recognize different cells in terms of cell types and status. The Transformer was applied to estimate the probability of sepsis by considering the global connectivity between capsules.

#### Capsule Network

The capsule network is a capsulating neural network architecture proposed for computer vision recognition [15]. In this study, we adopted the capsule network as a gene expression encoder due to its grouping architecture of genes into capsules through dynamic routing. The capsule network was aimed to group genes with similar biological functions into capsules that would represent immune related properties in sepsis.

The capsule network architecture for gene expression in this study contained two layers. The first layer was a fully connected layer that converted gene expression *X* = (*x*_1_,*x*_2_,…,*x*_*n*_) into eight primary capsules *u*_1_,*u*_2_,…,*u*_8_ by using different weights *W*_1_,*W*_2_,…,*W*_8_, where *u*_*i*_ = *W*_*i*_*X*. The second layer was the capsule layer that included 20 capsules *v*_1_,*v*_2_,…,*v*_20_, where each capsule was a vector with 16 dimensions calculated by dynamic routing. The primary capsules *u*_1_,*u*_2_,…,*u*_8_ were mapped to higher-level vectors 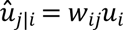 where *w*_*ij*_ is the weight matrix. Dynamic routing introduced a coupling coefficient 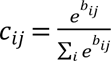 to concentrate on the important information within the primary capsule of genes without losing other features like maxpooling. The vector *s*_*j*_ was then composed of the 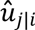 with coupling coefficient, which is 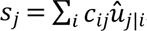. The capsule *v*_*j*_ was obtained by squashing, which was

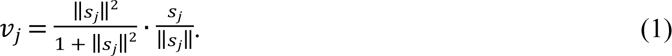

The first component of equation (1), 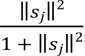, scaled the result to be within 0 and 1. The second component, 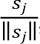, normalized the result and preserved the direction of the vector *s*_*j*_. The coupling coefficient *c*_*ij*_ included a coefficient *b*_*ij*_ which was updated as follows:

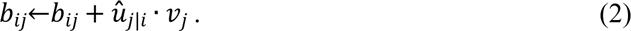

The coefficients in dynamic routing were updated by iteration in the following steps:

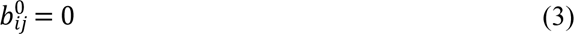

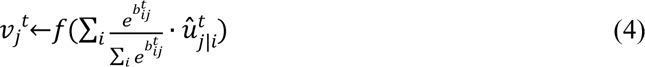

where 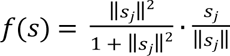.

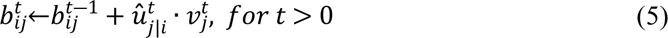

The dynamic routing was iterated for three times and the 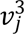 was the final capsule output. The network weight *w*_*ij*_ was obtained by backward propagation. The capsule network was applied as an encoder that represents different biological pathways through learning gene expression for septic classification.

#### Transformer

Transformer is a network architecture widely used in artificial intelligence, including natural language processing, computer vision, and content generation [16]. Through self-attention techniques, it can capture context by tracking relationships between distant elements in a sequence, making it well-suited for learning connectivity between different gene expression modules represented by capsules.

In this study, we used a multi-head attention mechanism as a decoder to decode the capsule output from the capsule encoder, denoted as

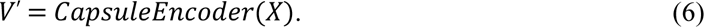

Each capsule output was transformed into a query matrix (Q), a key matrix (K), and a value matrix (V) by linear matrix transformations, i.e. *Q* = *W*_*q*_*V*^′^, *K* = *W*_*k*_*V*^′^, *V* = *W*_*v*_*V*^′^. The attention of the capsule output was then computed as follows:

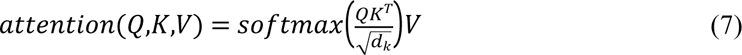

where *d*_*k*_ represented the dimension of the capsule, which was set to 16 in this study.

The output of a single self-attention head was denoted as *Z*_*i*_ = *attention*(*Q*_*i*_,*K*_*i*_,*V*_*i*_). We used multiple self-attention heads *Z*_1_,*Z*_2_,…,*Z*_*n*_, and concatenated them to *Z*^′^. Finally, we applied a dense layer with 150 neurons and a sigmoid activation function to obtain the probability of sepsis occurrence based on the concatenated attention output.

### Pre-Training on Single-Cell RNA-Seq

The cells collected from 29 septic patients and 36 corresponding controls were normalized and integrated. They were then randomly divided into three sets: the training set (80% of the cells), the validation set (10% of the cells) to fine tune the neural network, and the test set (10% of the cells) to evaluate the performance of the scCaT.

To optimize the output probability, we applied binary cross-entropy loss function:

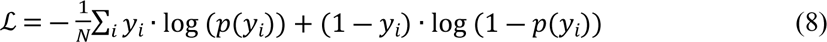

where *y*_*i*_ is the label of the cells, and *p*(*y*_*i*_) is the predicted probability obtained using scCaT *ŷ*_*i*_ = *Transformer*(*CapsuleEncoder*(*x*_*i*_)). We performed backpropagation to update the network parameters using Adam optimization based on the loss function. The network was trained for 50 epochs with a batch size of 32 and early stopping.

After training, the network model was tested on the test set, which consisted of 4,265 cells, based on area under ROC curve (AUC). The AUC shows the trade-off between true positive rate (TPR) and false positive rate (FPR) that were calculated as follows:

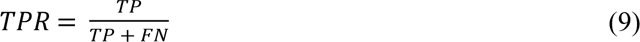

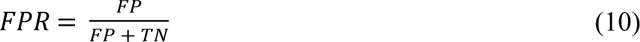

where TP is the number of true positively classified samples, FN is the number of false negatively classified samples, FP is the number of false positively classified samples, and TN is the number of true negatively classified samples.

We compared the performance of scCaT to that of existing biomarkers including FAIM3/PLAC8, SeptiCyte, and sNIP, and the traditional machine learning methods, including decision tree, random forest, naïve bayes, K-nearest neighborhoods, and quadratic discriminant analysis on the single-cell RNA-seq test set.

### Transferring To Bulk RNA Cohorts

Our final model aimed to assist diagnosis in the clinic, so we attempted to deploy it on a different data type using transfer learning. We utilized the scCaT trained on single-cell RNA-seq as a pretrained model and fine-tuned it on bulk RNA-seq and microarray data. By following the same procedure, our model could be transferred to other types of data commonly used in the clinic.

We collected nine microarray cohorts and one bulk RNA-seq for transfer and evaluation (**Table 1**). The three largest microarray cohorts, GSE95233, GSE26440, and GSE57065, were integrated to fine-tune the pretrained network. We initialized the last dense layer and froze all the previous layers, including the capsule and self-attention layers. In this way, we trained the last layer on the three largest cohorts and then trained the entire network to focus more on microarray data. Following the same procedure, we fine-tuned the pretrained network on 30% samples in bulk RNA-seq cohort, GSE185263, which contained 103 sepsis patients and 14 healthy controls.

We evaluated the performance of the fine-tuned network on six independent cohorts and 70% samples in the bulk RNA-seq cohort, GSE185263, using AUROC and compared it to existing biomarkers, including FAIM3/PLAC8, SeptiCyte, and sNIP, as well as traditional machine learning methods such as decision tree, random forest, naïve bayes, K-nearest neighborhoods, and quadratic discriminant analysis. We also conducted a rotated test, where we transferred the model to one cohort and tested it on other cohorts.

### Annotation of Network Model

To investigate the biological functions learned by scCaT from gene expression, we extracted the capsule outputs and performed visualization using uniform manifold approximation and projection (UMAP) with cell type annotation.

To resolve the function of the capsule network, we visualized and extracted the genes captured in the primary capsule layer. We extracted the genes’ weights for each of the eight primary capsules and displayed the most important genes in a heatmap. Next, we compared the genes with higher importance, whose weights were larger than 0.06 in absolute value, across the eight primary capsules. To gain insight into the biological pathways associated with each capsule, we performed enrichment analysis of the important genes using Gene Ontology (GO).

To uncover the pathways of capsules, we conducted an activation test on the capsule outputs and retrieved genes participating in each capsule. Specifically, we systematically inhibited some of the gene inputs to determine which genes made large positive or negative contribution to each capsule. For instance, the capsule 8 and capsule 19 were significantly differential when they had particular genes activating. We then utilized Gene Ontology (GO) to investigate the biological pathways associated with the activated genes, which enabled us to infer the capsule pathways learned by the capsule network for each of these capsules. Subsequently, we constructed a capsule-pathway network to better understand the relationships between the biological functions of the 20 capsules.

## Results

### Overall Framework

scCaT is a novel framework that incorporates the capsulating architecture with Transformer and the transfer learning from single-cell RNA sequencing (scRNA-seq) data to bulk RNA data. We collected cells from patients with sepsis and cells from control individuals (**Figure 1A**). Then, we used 80% of the cells as training samples to train the capsulating architecture with Transformer (**Figure 1B**), while the remaining cells were used for tuning hyperparameters and single-cell validation. The capsulating architecture is also referred as capsule network, where genes adopted in capsules was updated by dynamic routing (**Figure 1C**). The model trained on scRNA-seq was treated as a pretrained model, and we then transferred it to bulk RNA data, specifically microarray and bulk RNA-seq data. The neural network was then fine-tuned on the three largest cohorts out of nine microarray profiles and a small proportion of a bulk RNA-seq profile, while the remaining six cohorts and the rest of the bulk RNA-seq profile were used for validation and comparison to other biomarkers and traditional machine learning methods (**Figure 1D**).

**Figure 1.**
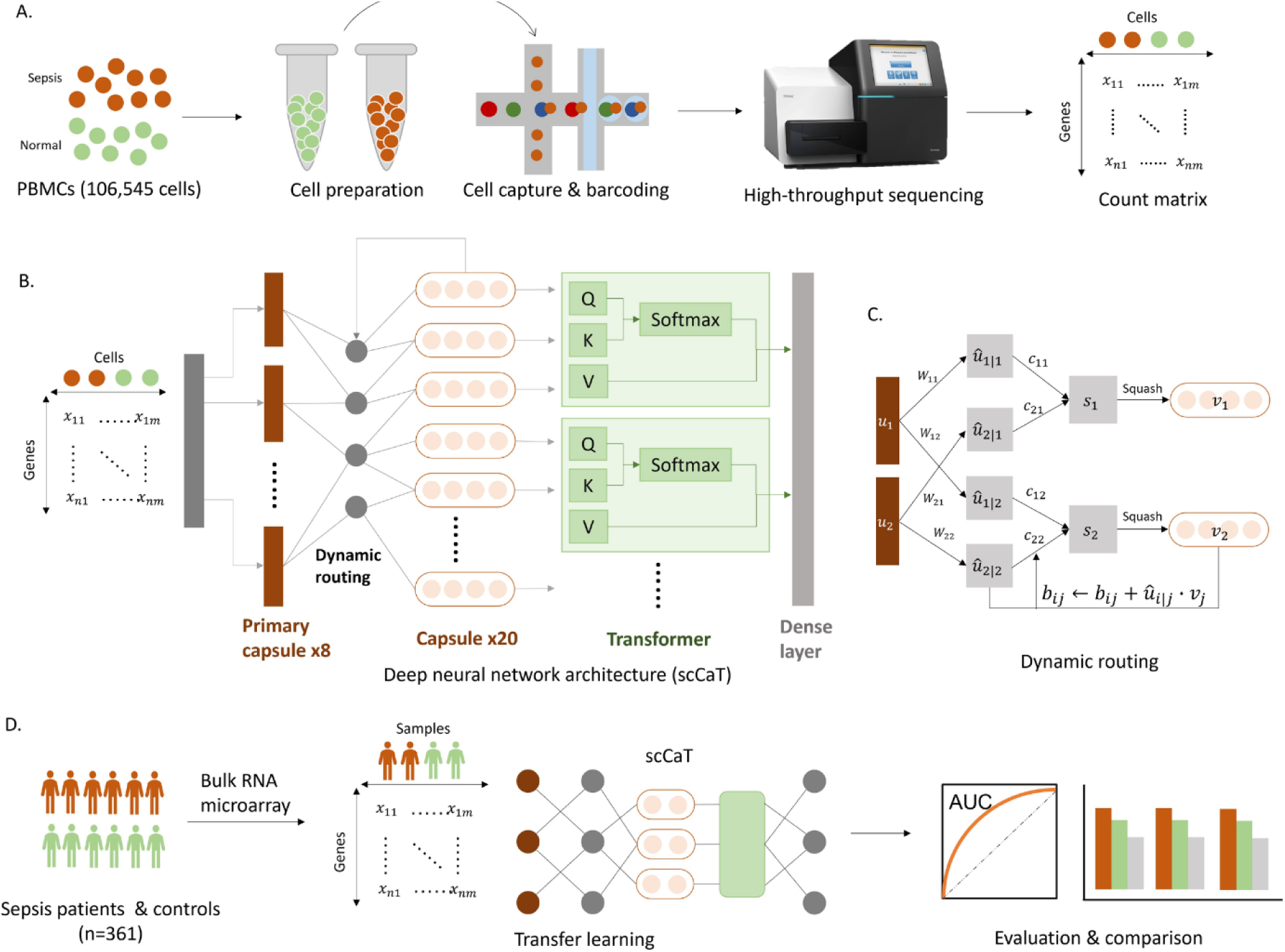
Study workflow diagram. A. Single-cell gene expression of peripheral blood mononuclear cells collected from sepsis patients and normal controls. B. Deep neural network architecture of scCaT. scCaT was constructed by blending capsule network and Transformer, and then it was trained using the gene expression of cells. C. Dynamic routing procedures of scCaT. D. Transfer learning. scCaT was transferred to subjects using bulk RNA data for fine-tune. It was evaluated and compared on independent cohorts.

### Performance Evaluation on Single-Cell RNA Sequencing Data

In order to assess the performance of scCaT, we conducted a test on single-cell RNA sequencing data’s test set, which accounted for 10% of the SCP548 dataset. We compared scCaT with existing biomarkers, such as FAIM3/PLAC8 [14], SeptiCyte [13], and sNIP [12], as well as traditional machine learning models like nearest neighbors, decision tree, random forest, naïve bayes, and quadratic discriminant analysis.

scCaT achieved an AUROC of 0.93, which is higher than FAIM3/PLAC8 (0.49), SeptiCyte (0.60), and sNIP (0.51) (**Figure 2A**). The nearest neighbors, decision tree, random forest, naive bayes, and quadratic discriminant analysis models achieved AUROC scores of 0.61, 0.71, 0.87, 0.68, and 0.77, respectively. The superior classification capability of scCaT mainly attributed to the higher complexity and parameter count for scRNA-seq data that contained different cell types (as explained below).

**Figure 2.**
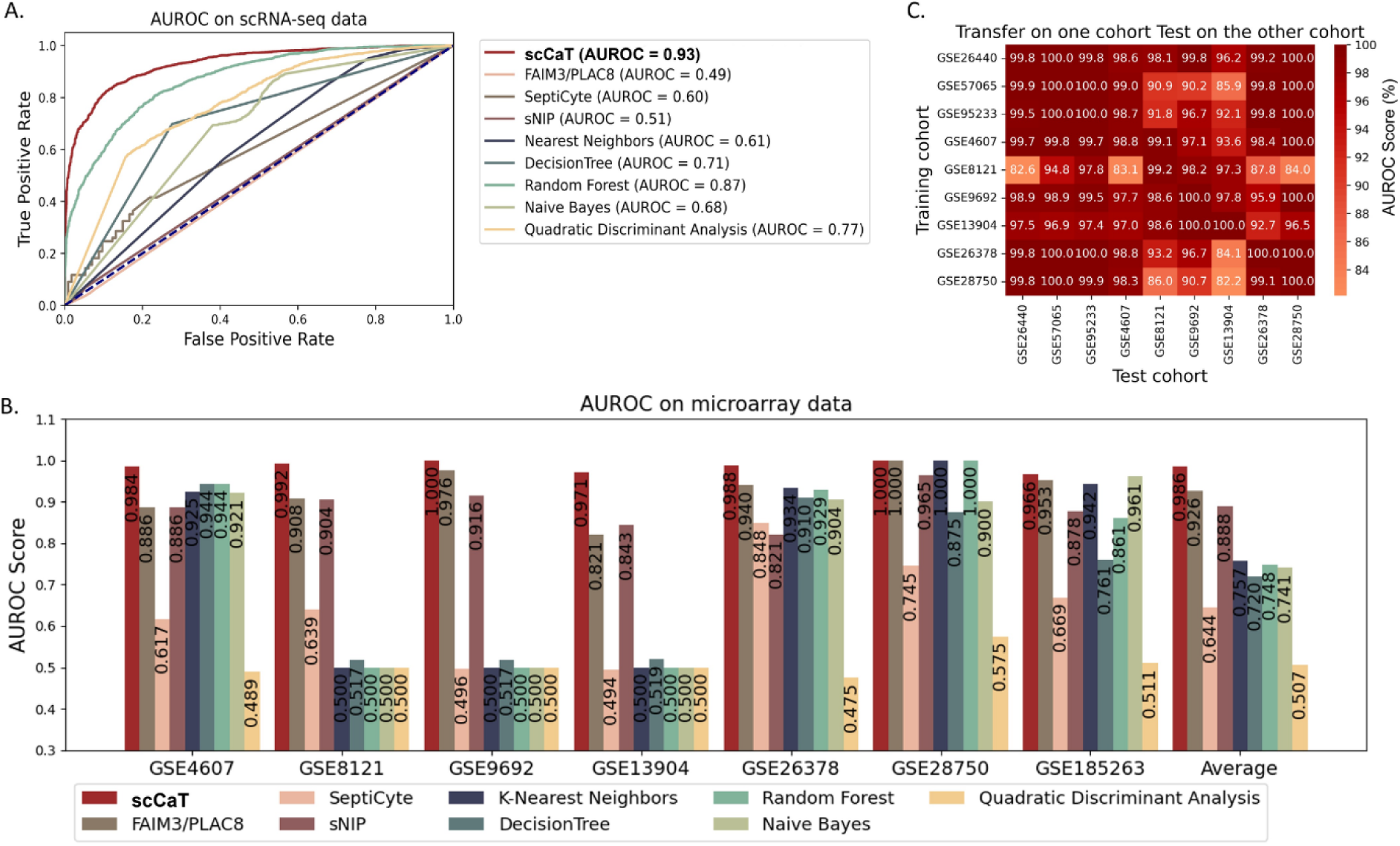
Performance of scCaT in scRNA-seq data and microarray cohorts. A. ROC curve demonstrating the performance of scCaT, existing biomarkers, and traditional machine learning methods, for sepsis diagnosis from single-cell data. B. AUROC score demonstrating the performance of scCaT and the other methods for sepsis diagnosis from microarray data. C. Heatmap showing the AUROC score of scCaT transferred on one cohort and tested on the others.

### Performance Evaluation on Bulk RNA Data

To transfer scCaT on other data type, we applied the pre-trained model on scRNA-seq data and fine-tuned on another data type. Specifically, we fine-tuned the model on the three largest microarray cohorts and evaluated its performance on six independent validation cohorts. We also fine-tuned the model on a small proportion of bulk RNA-seq samples and evaluated it on the rest 70% samples of the bulk RNA-seq cohort.

We compared the scCaT’s performance to that of existing biomarkers (FAIM3/PLAC8, SeptiCyte, and sNIP) and the traditional machine learning methods (nearest neighbors, decision tree, random forest, naive Bayes, and quadratic discriminant analysis) on these six microarray cohorts and one bulk RNA-seq cohort. ScCaT outperformed all other methods with an average AUROC of 0.971∼1.000 (**Figure 2B**). Although FAIM3/PLAC8 achieved comparable performance to our model on average, it performed poorly on GSE4607 and GSE13904 (AUROC of 0.886 and 0.821 respectively) compared to our model (AUROC of 0.984 and 0.971, respectively).

Notably, FAIM3/PLAC8 and sNIP were discovered from microarray data, which may explain their comparable performance to our model on microarray. However, these biomarkers lack the robustness to be transferred to other data types for clinical use. In contrast, our framework allows for fine-tuning of the model, making it applicable in a clinical setting with only a few clinical-type data.

To assess the generalization performance of the model, we conducted a rotated test on the nine microarray cohorts, where the model was fine-tuned on one cohort and tested on another one (**Figure 2C**). The results demonstrate that scCaT achieved an AUROC above 0.90 in most cases, with the worst case having an AUROC of 0.82. These findings indicate that our framework has good generalization performance.

### Cell Types Learned by Capsule Network

The capsule network played an important role in the model by adopting the associated genes into capsules and providing encoder for the whole model. We visualized the capsule outputs using UMAP and found that the capsule network can learn the cell type automatically (**Figure 3**). Compared to the raw input single-cell data (**Figure 3A**), the capsule network transformed the genes into capsules and clustered the cells into distinct groups (**Figure 3B**). Interestingly, within each cluster, the cells from sepsis and control samples were separated.

**Figure 3.**
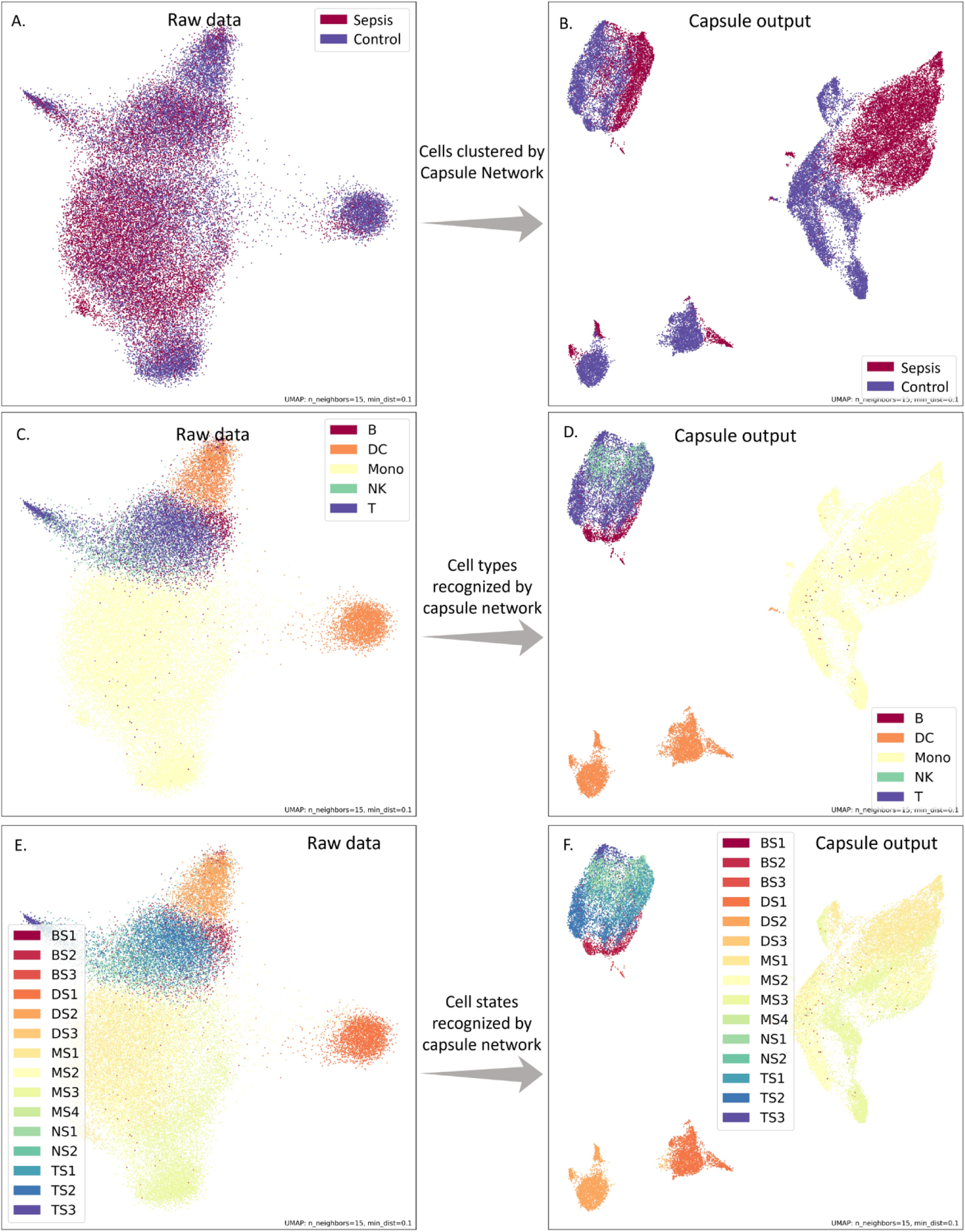
UMAP plot demonstrating the status, cell types, and cell states generated by raw data and capsule outputs. The raw input data (A) and the capsule outputs (B) annotated by samples collected from sepsis patients and controls. The capsule outputs clustered cells in several groups and separate sepsis and controls in each group. By annotating the raw input data (C & E) and the capsule outputs (D & F) with cell types and cell states, we found that the group clustered by the capsule outputs corresponded to different cell types and cell states.

To further investigate the capabilities of the capsule network, we annotated the cells by their cell types and states. We found that the capsule outputs clustered different cell types and cell states in comparison to the raw input (**Figure 3C-F**), indicating that the capsule network can learn cell type information. The capsule network can group B cells, T cells, and natural killer cells, which are differentiated from lymphoid progenitor, into the same cluster (**Figure 3D**). Moreover, the capsule network was able to separate two dendritic cell states (DS1 and DS2 in **Figure 3F**).

Our observations suggested that the differences across cell types were greater than the differences between cells from sepsis and controls. Theoretically, the capsule network first learned how to distinguish different cell types and then classified cases and controls within each cell type. Besides that, we found that each dimension of the capsules can be used to represent an aspect of the capsule outputs (**Supplementary Figure S2** and **S3)**.

### Functional Characterization of Primary Capsules

To reveal the genes participating in primary capsules, we extracted and visualized their weights for each gene (**Figure 4A**). We identified the genes with higher importance and analyzed their intersection between primary capsules using an upset graph (**Figure 4B**). The results showed that only a few genes were shared among two or three primary capsules, indicating that each primary capsule captured distinct gene information from different aspects.

**Figure 4.**
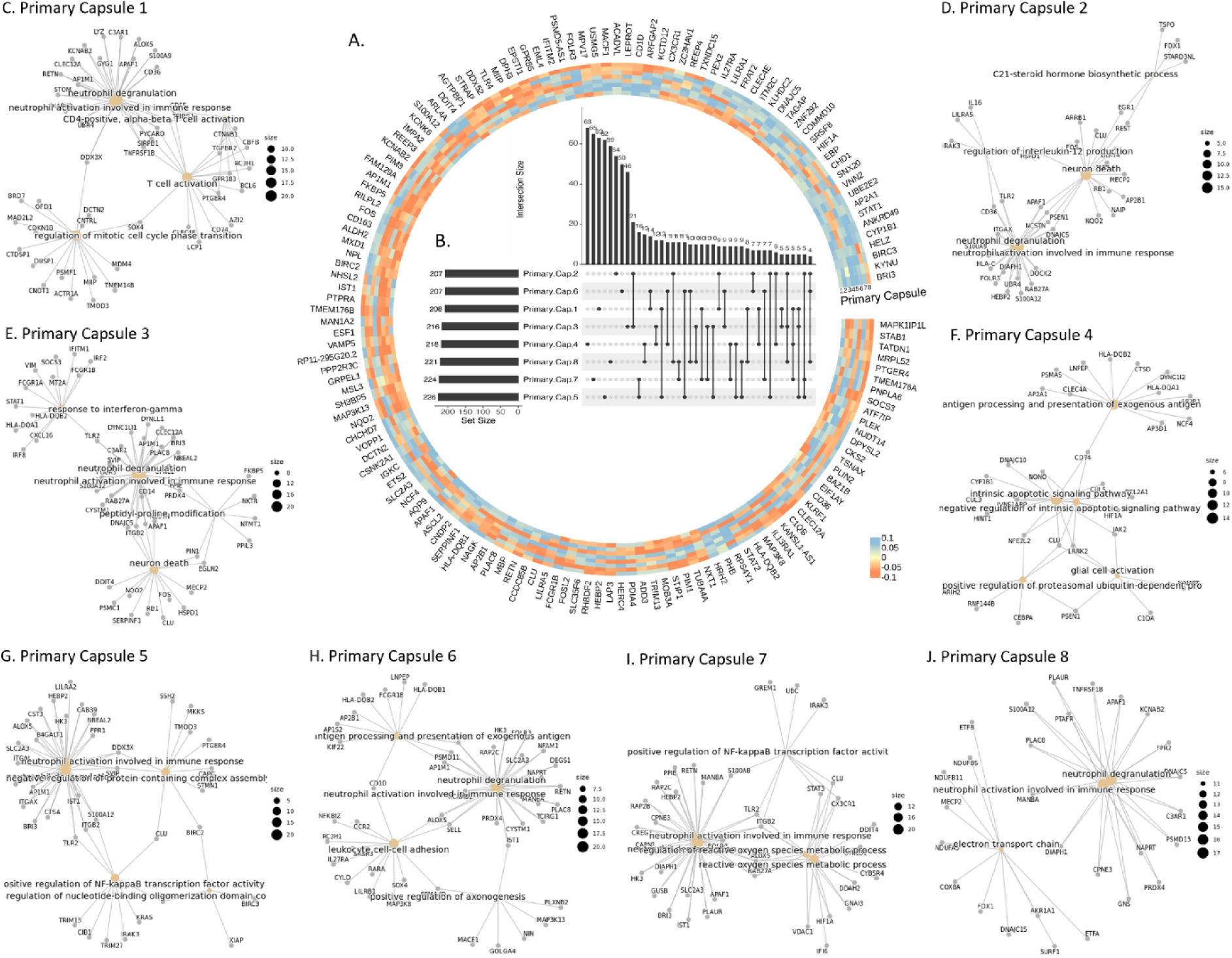
Explanation of the primary capsule layer in scCaT. A. Heatmap visualization of the weights of the eight primary capsules. B. Upset graph showing the intersection of genes included in the eight primary capsules. C-J. Pathway analysis of the genes included in each of the eight primary capsules. The enriched biological pathways are different from the other primary capsules. Gray node represents genes included in each capsule and yellow node represents enriched pathway.

We further explored the biological pathways associated with the genes in primary capsules. Except for primary capsule 4, the genes extracted by the primary capsules were mainly involved in neutrophils degranulation and activation, which are critical events in sepsis pathogenesis and can cause tissue damage [17] (**Figure 4C-J**). Moreover, each primary capsule was also associated with specific pathways related to inflammation in sepsis. For instance, primary capsule 5 and 7 were involved in the NF-kappaB transcription factor activity, a central mediator of pro-inflammatory gene induction and functions [18] (**Figure 4G**). The primary capsule 2 was involved in neuron death and the regulation of interleukin-12 production, a proinflammatory cytokine with immunoregulatory functions [19] (**Figure 4D**). Other primary capsules were also involved in pathways related to the inflammation in sepsis, such as primary capsule 1 for T-cell activation (**Figure 4C**), primary capsule 3 for neuron death and interferon-gamma [20] (**Figure 4E**), primary capsule 4 for apoptotic signaling pathway (**Figure 4F**), primary capsule 6 for leukocyte cell-cell adhesion [21] (**Figure 4H**), and primary capsule 7 for reactive oxygen species metabolic process [22] (**Figure 4I**).

### Capsule-Pathway Network Analysis

After the primary capsule layer, genes were further grouped into several capsules based on their biological functions by dynamic routing. To identify the genes in each capsule, we conducted activation analysis in the capsule network. Specifically, we selectively activated or shut down certain gene inputs and identified the genes that contributed most to each capsule through enumeration (**Figure 5A** and **5B**). These important genes were learned by the capsule network and helped classify sepsis and control samples.

**Figure 5.**
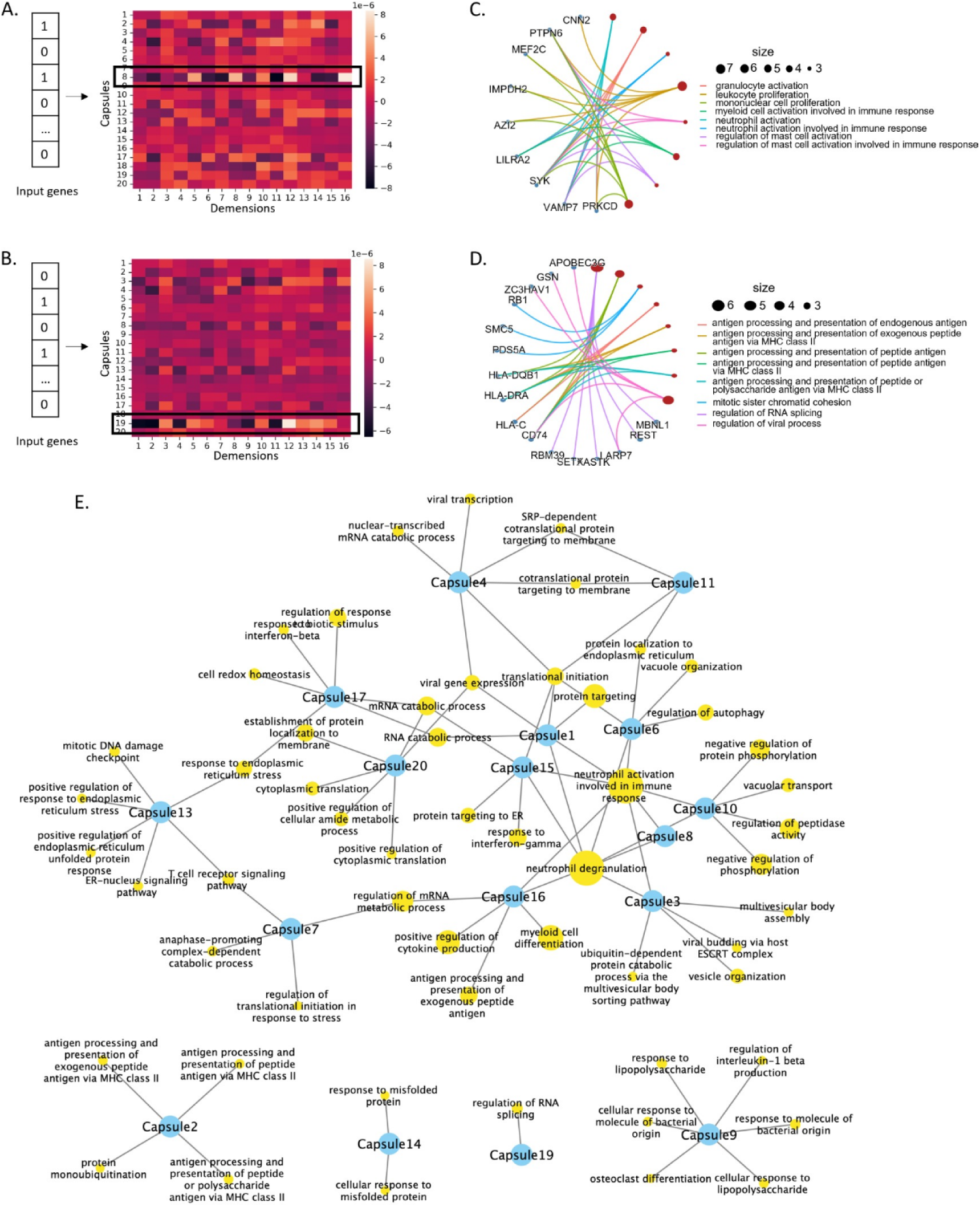
Capsule pathways learned by scCaT. A-B. By activating specific patterns of genes as input, the capsule was activated and the specific genes adopted in each capsule can be identified. C-D. The biological pathways enriched in capsules 8 and 19. E. Capsule-pathway network. Blue nodes represent capsules and yellow nodes represent pathways.

Next, we identified the capsule pathways, i.e., the functions enriched in each capsule, based on Gene Ontology (GO) (**Supplementary Figure S1**). For instance, the pathways of capsule 8 included neutrophil activation and neutrophil degranulation that are critical in inflammation, the proliferation of lymphocyte, nonnuclear cell, and leukocyte, which are hallmark of the adaptive immune response to pathogens [23] (**Figure 5C**). The capsule 19 was involved in RNA splicing, regulation of viral process, cellular response to interferon-gamma, and the antigen processing, indicating that it grouped genes related to the immune responses to viral infection (**Figure 5D**).

Finally, we constructed a capsule-pathway network of the 20 capsules. Only a few genes were shared between different capsules. Our results suggested that genes in each capsule were mainly enriched in several pathways, indicating that the capsules filtered genes based on different biological aspects. We also found that Capsules 1, 3, 6, 8, 10, and 15 participated in the activation and degranulation of neutrophils, which are critical in the inflammation seen in sepsis (**Figure 5E**). Furthermore, different capsules enriched in specific biological functions, suggesting that they can learn about cells and sepsis by considering different functions from different perspectives.

For example, Capsule 9 in the capsule-pathways network focuses on the a series of bacterial infection-related functions different from other capsules, including response to molecule of bacterial origin, response to lipopolysaccharide, regulation of interleukin-1 beta production and osteoclast differentiation (**Figure 5E**). Lipopolysaccharide is produced by Gram-negative bacteria and can activate innate immune responses to the molecule of bacterial origin [24]. The bacteria-induced inflammation is associated with overproduction of cytokines including interleukin-1, which can amplify osteoclast differentiation [25].

## Discussion

We developed scCaT, a deep learning framework that combines capsule network and Transformer to create an explainable model for sepsis detection based on gene expression. We identified the genes involved in each capsule and discovered that scCaT’s capsulating architecture groups genes with similar biological pathways into 20 capsules, which can differentiate between various cell types and distinguish sepsis from control samples in each cell type.

The capsules in this study were acquired using dynamic routing instead of backward propagation. Dynamic routing involved updating the coefficient *c*_*ij*_ of each primary capsule in a way that each capsule output would concentrate more on one or several primary capsules. This was achieved through the update of *b*_*ij*_ using Equation (2). If the capsule output was similar to one of the primary capsules, the *b*_*ij*_ would assign greater weight to that primary capsule through iterations. The number of iterations determined the strength of this weight. In our study, we utilized three times of iterations since an excessive number of iterations could eliminate dissimilar primary capsules and render the process redundant.

It is of vital importance for researchers to gain insights into the specific genes, pathways, or biological processes that contribute to pathogenesis, enabling further exploration and refinement of diagnostic criteria. Activation tests on the capsule explicitly show that the genes adopted in different capsules have only a few intersections, indicating that the capsules extracted features from different aspects (**Figure 4**). Each capsule possesses specific biological functions related to sepsis and identifies sepsis based on these functions. Gene ontology analysis show that each capsule enriched in particular functions related to different processes in inflammation (**Figure 5**).

In the capsule-pathway network, it seems that capsules focus on a specific series of pathways regulated in sepsis inflammation such as Capsule 9 (**Figure 5E**). Moreover, some capsules focused on biological pathways such as the degranulation and activation of neutrophils, indicating that the capsule network has learned the critical events in sepsis pathogenesis that can cause tissue damage. Furthermore, different capsules enriched in specific biological functions, suggesting that they can learn about cells and sepsis by considering different functions from different perspectives.

The model was pretrained on single-cell RNA-seq data and then transferred to bulk RNA-seq and microarray cohorts using transfer learning. Following the same procedure, scCaT can also be adapted to the data types used in clinical practice, such as RT-PCR. Transfer learning has two advantages: (1) deploying the model across data measured by different biological techniques, and (2) improving the performance by associating large datasets to pretrain the model. This study provides a framework for transferring deep learning model to clinical settings and diagnosing rare diseases with insufficient data for training.

Although scCaT has exceptional performance, there is room for improvement. As the omics data in the sepsis research field is accumulating rapidly, in our future work, we plan to enhance scCaT by integrating diverse sources of datasets, including genomics, proteomics, and metabolomics. Furthermore, we will keep collecting clinical data and sepsis samples to fine-tune and attempt to deploy the model in clinical applications after clinical validation.

The study demonstrated a method for learning gene modules with a capsulating architecture and transferring the model to other data types. This framework could be useful in diagnosing rare diseases where obtaining a large number of subjects is challenging.

## Authors’ Contributions

X.Z. and L.C. conceived the idea and drafted the manuscript. X.Z., D.M., and D.C. implemented the idea and prepared figure 1-5. W.W. and K.T. conducted the initial experiments. D.M., J.W., and L.Z. gave suggestions and helped write the manuscript. Y.L. prepared figure 5C and 5D. K.L., M.W., and L.C. supervised the project. All authors reviewed the manuscript.

## Consent For Publication

Not applicable.

## Availability Of Data And Materials

The source code of scCaT has been uploaded to https://github.com/Kimxbzheng/CaT. The data cohorts used in this study can be found on Broad Institute Single Cell Portal (https://singlecell.broadinstitute.org/single_cell), with portal ID: SCP548 and Gene Expression Omnibus (GEO) database according to their accession ID.

